# A deep learning pipeline for time-lapse camera monitoring of insects and their floral environments

**DOI:** 10.1101/2024.04.12.589205

**Authors:** Kim Bjerge, Henrik Karstoft, Hjalte M. R. Mann, Toke T. Høye

## Abstract

Arthropods, including insects, represent the most diverse group and contribute significantly to animal biomass. Automatic monitoring of insects and other arthropods enables quick and efficient observation and management of ecologically and economically important targets such as pollinators, natural enemies, disease vectors, and agricultural pests. The integration of cameras and computer vision facilitates innovative monitoring approaches for agriculture, ecology, entomology, evolution, and biodiversity. However, studying insects and their interactions with flowers and vegetation in natural environments remains challenging, even with automated camera monitoring.

This paper presents a comprehensive methodology to monitor abundance and diversity of arthropods in the wild and to quantify floral cover as a key resource. We apply the methods across more than 10 million images recorded over two years using 48 insect camera traps placed in three main habitat types. The cameras monitor arthropods, including insect visits, on a specific mix of *Sedum* plant species with white, yellow and red/pink colored of flowers. The proposed deep-learning pipeline estimates flower cover and detects and classifies arthropod taxa from time-lapse recordings. However, the flower cover serves only as an estimate to correlate insect activity with the flowering plants. Color and semantic segmentation with DeepLabv3 are combined to estimate the percent cover of flowers of different colors. Arthropod detection incorporates motion-informed enhanced images and object detection with You-Only-Look-Once (YOLO), followed by filtering stationary objects to minimize double counting of non-moving animals and erroneous background detections. This filtering approach has been demonstrated to significantly decrease the incidence of false positives, since arthropods, occur in less than 3% of the captured images.

The final step involves grouping arthropods into 19 taxonomic classes. Seven state-of-the-art models were trained and validated, achieving *F*1-scores ranging from 0.81 to 0.89 in classification of arthropods. Among these, the final selected model, EfficientNetB4, achieved an 80% average precision on randomly selected samples when applied to the complete pipeline, which includes detection, filtering, and classification of arthropod images collected in 2021. As expected during the beginning and end of the season, reduced flower cover correlates with a noticeable drop in arthropod detections. The proposed method offers a cost-effective approach to monitoring diverse arthropod taxa and flower cover in natural environments using time-lapse camera recordings.

## 1. Introduction

Arthropods including insects, constituting the most diverse group contributing to 50% of the animal biomass (Bar-On et al., 2018), play vital roles in terrestrial ecosystems and hold significant economic importance, serving as both agricultural pests and pollinators. Conventional insect trapping techniques, as outlined in Montgomery et al. (2021), involve the manual collection and sampling of insects. Invariably, insects are also sacrificed and the work of manual enumeration and taxonomic identification by human experts necessitates specialised knowledge and is very labour intensive. Moreover, this method poses a threat to rare insect species due to the fatal nature of the trapping process.

The advent of automated monitoring technologies, employing computer vision and deep learning, has brought about a revolution to insect surveys (van Klink et al., 2022; Lima et al., 2020; Besson et al., 2022). Computer vision methods, demonstrating substantial promise in both real-time scenarios (Bjerge et al., 2021a,b) and offline analysis of images from time-lapse (TL) cameras (Geissmann et al., 2022; Bjerge et al., 2023a), have significantly advanced insect monitoring capabilities. Automated insect camera traps, coupled with data-analyzing algorithms rooted in computer vision and deep learning, serve as invaluable tools for comprehensively monitoring insect trends and elucidating the underlying drivers (Barlow and O’Neill, 2020; Høye et al., 2021).

Automation is crucial for achieving spatiotemporal upscaling of global insect monitoring and facilitating efficient pest management (Preti et al., 2021). The increasing utilization of deep-learning methods for classification in the realm of entomology (Høye et al., 2021), underscores the need for large training datasets to ensure robust predictions. However, the diverse nature of insects, coupled with highly variable abundance, often results in unbalanced datasets (Werner de Vargas et al., 2023) with numerous classes, posing a limiting factor in the application of deep learning to entomological contexts. As these methods mature and become integral components of monitoring programs, the demand for robust techniques to detect, filter and classify species in natural environments becomes increasingly important.

The majority of insects, particularly pollinators, are attracted to flowering plants, making it crucial to monitor them in conjunction with arthropod abundance. Predators, like spiders, are also presumed to be indirectly attracted to flowers due to the expected increase in insect prey during the flowering season. The flower coverage observed from the camera perspective is expected to influence pollinators and should be adjusted to correlate with measurements of insect abundance and activity. Hence, it is crucial to include an estimate of flower cover in the recorded images as part of the image processing pipeline.

### 1.1. Related work

The automation of arthropod detection poses a substantial challenge due to the swift movement of insects, the ephemeral nature of their environmental interactions (such as pollination events) and the diminutive sizes of the animals (Høye et al., 2021; Xia et al., 2018). Furthermore, insects and other arthropods may be occluded by flowers or leaves, complicating the separation of objects of interest from the surrounding natural vegetation. An object in an image that resembles an insect for the model, e.g. a small part of the vegetation, may lead to multiple false detections in consecutive images Bjerge et al. (2021b).

Images captured by fixed-position cameras may not capture distinguishing features or provide sufficient resolution for confident species identification, particularly for those with similar morphologies. Typically, the working distance of the camera to the floral scene is in the area of 50 cm which means that arthropods constitute only a small portion of each image, often lacking the details necessary for precise species identification. Therefore, it is essential to filter false positive detections and categorise arthropods into taxonomic groups based on the image information content captured in the current study.

Convolutional neural networks (CNNs) have emerged as a powerful tool in deep learning for the localization and classification of insects and various organisms. Existing research has explored insect localization and classification through CNNs in diverse contexts (Hansen et al., 2020; Kasinathan et al., 2021; Kittichai et al., 2021; Xia et al., 2018), addressing tasks ranging from categorizing museum specimens to detecting insects in field crops like rice, soybeans and other agricultural settings. Notably, Bjerge et al. (2023a) achieved high accuracy in insect localization and classification, albeit in a study limited to a specific location with low biodiversity and a restricted number of species. Research has also delved into classifying insects into taxonomic ranks, such as order, family and genus (Ong and Hamid, 2022; Xia et al., 2018; Bjerge et al., 2023c). This approach proves particularly relevant for monitoring sites characterised by high species diversity. However, the challenge persists in collecting sufficient training samples to encompass the vast array of species within diverse ecosystems.

While previous studies have proposed insect detection and tracking methods in real-time and offline settings (Bjerge et al., 2021a; Ratnayake et al., 2021), these works primarily focus on continuous video recordings, posing challenges for long-term, in-field camera recordings. Such continuous recording requires substantial hardware capabilities for data processing and storage compared to time-lapse camera trapping. Surprisingly, none of the aforementioned studies address the combined monitoring of flowers with arthropod detection and classification. Conversely, various publications have focused on flower and plant segmentation (Nilsback and Zisserman, 2010; Aydin and Uǧgur, 2011; Al-Vshakarji et al., 2017; Inthiyaz et al., 2018; Lu et al., 2022). Many of these proposals utilise traditional computer vision methods, such as colored texture feature representation, color clustering, histogram-based thresholding, watershed segmentation and foreground/background segmentation using shape measurements. However, most methods concentrate on segmenting individual leaves and flowers in datasets with clearly defined, visible, single, or few flowers. This emphasis on well-defined backgrounds may not effectively address the diversity and complexity of backgrounds encountered in monitoring arthropods interacting with flowers in natural environments.

### 1.2. Contribution

Our paper tackles the intricate task of estimating flower cover and identifying and counting arthropods in insect camera trap time-lapse sequences, which may become more abundant in the future. However, the study is limited to a particular composition of *Sedum* plants which enables a uniform method to attract pollinators on different monitoring locations. We introduce an extensive image processing pipeline that harnesses the power of deep learning techniques to monitor insects and flowers through time-lapse camera recordings in natural settings. This pipeline integrates approaches for arthropod detection, classification and flower cover estimation, providing an effective solution for biodiversity monitoring. The efficacy of our approach is demonstrated through rigorous evaluation on a substantial dataset. To our knowledge, no other publications have addressed the monitoring of insect populations in flowering environments on the temporal and spatial scale as presented in our work. Our primary objectives for analysing time-lapse camera recordings are as follows.

- Propose a deep learning pipeline to measure temporal abundance for selected taxa of arthropods
- Filter false positive background detections and remove double counting of non-moving arthropods
- Estimate the flower cover of the vegetation for studies of season dynamics for both flowers and insects
- Demonstrate the proposed method on camera recordings for a whole monitoring season

## 2. Method

This work aims to introduce a comprehensive pipeline shown in Figure 1 for the estimation of flower cover, detecting and classifying arthropods in image recordings obtained from a natural environment. The images were captured using time-lapse cameras specifically chosen for monitoring the vegetation including a mix of 12 species of *Sedum* plants and their associated arthropod activity. The deliberate selection of *Sedum* plants is driven by their low vegetation, which facilitates effective camera focus on both flowering plants and visiting arthropods, encompassing insects and pollinators.

**Figure 1:**
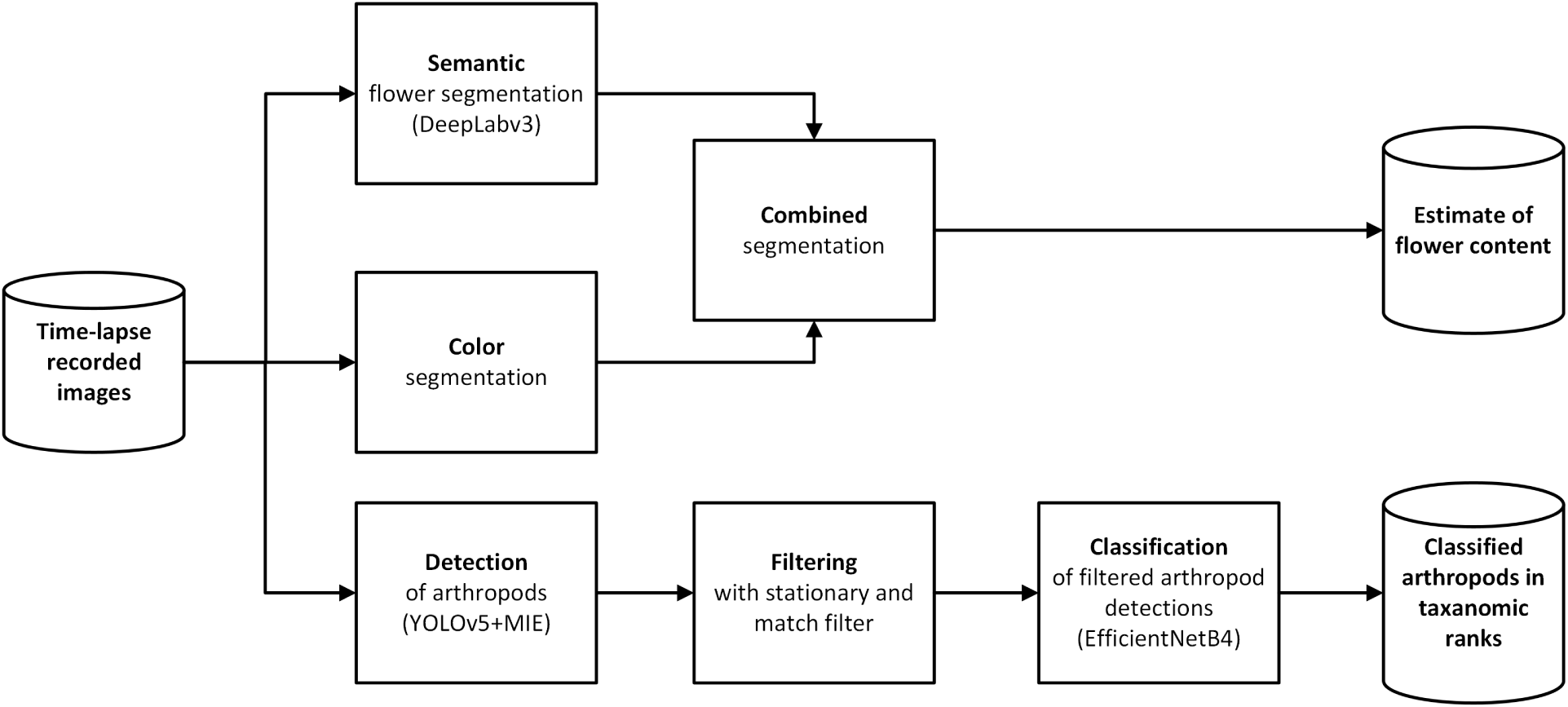
This figure illustrates the processing pipeline used to estimate flower cover and classify arthropod taxa in time-lapse image recordings. Flower cover is estimated by applying a logical AND operation to image pixels of semantic and color-segmented images. Arthropod detections are filtered by removing non-moving objects using a stationary and match filter. This filter reduces false positive detections and double counting of non-moving insects. The filtered arthropod detections are then classified into 19 taxonomic categories, including plant parts and undefined species.

### 2.1. Estimating flower cover in images

Numerous arthropods exhibit an attraction to flowering plants, particularly insect pollinators such as Hymenoptera and Lepidoptera, encompassing honeybees, bumblebees, hoverflies and butterflies. Consequently, we predict that the cover of flowers is correlated with the occurrence of visiting insects and other arthropods and we employed a hybrid deep learning approach by quantifying both semantic and color segmentation. Semantic segmentation, as reviewed by Guo et al. (2018) and Yu et al. (2018), poses a challenging problem in computer vision. The objective is to assign a label to each pixel in an image, ensuring that pixels with the same label share specific characteristics of information within each region. Semantic segmentation, a subset of image segmentation, involves labelling every individual pixel in an image according to its semantic class or region. In our context, we classify each pixel as belonging to either background vegetation or flowers. This approach is simpler and more flexible than establishing a semantic class for each individual flower species.

The survey by Minaee et al. (2022) on image segmentation using deep learning identifies 11 categories of Deep Learning (DL) based image segmentations models. Among theses, Fully Convolutional Networks (Shelhamer et al., 2017) (FCNs) made a milestone in DL-based semantic image segmentation models. Encoderdecoder based models as U-net (Ronneberger et al., 2015) comprises two parts, a contracting path to capture context, and a symmetric expanding path that enables precise localization. The U-Net model is particularly effective at learning from very few annotated images through data augmentation. DeepLabv3 (Chen et al., 2018) builds on the encoder-decoder strategy by incorporating Atrous Spatial Pyramid Pooling (ASPP), a technique that applies multiple sampling rates to a convolutional feature layer. This approach allows

DeepLabv3 to capture objects at various scales and robustly segment them within a multiscale image context. Due to its ease of use and high performance, DeepLabv3 has become a widely popular semantic segmentation model, achieving an impressive 85.7% mIOU on the PASCAL VOC 2022 dataset, placing it among the best state-of-the-art models and therefore used in our work.

In semantic segmentation, an inherent challenge lies in accurately labelling regions. To address this challenge, we leveraged the color features of plants and flowers to automate the labelling process through the creation of a color mask for flowers. Initially, we generated a flower mask encompassing yellow, white and red/pink flowers. This involved converting the image into pixel values represented in the Hue, Saturation, Value (HSV) format. Samples of the three different colored flowers were utilised to determine a threshold for the HSV values, effectively separating flowers from plants. An example of threshold color mask is shown in Figure 2. However, color segmentation is susceptible to variations in illumination and changes in the background plant’s appearance, particularly during the later stages of the season when leaves may transition to brownish or reddish hues. Consequently, the dataset used for training and testing underwent manual sorting and verification, as detailed in Section 3.1.

**Figure 2:**
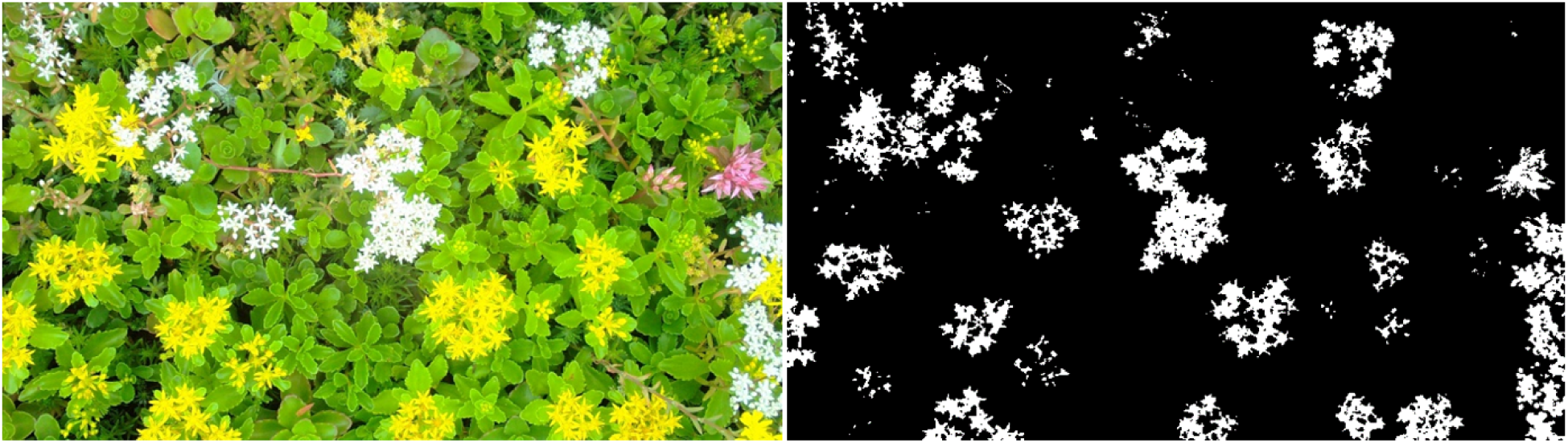
Example of image with flowers and a binary black and white flower mask

Subsequently, employing the dataset of flower masks, we trained a DeepLabv3 semantic segmentation model using transfer learning published by Minhas (2019). The trained model produces a segmented image, distinguishing between background plants and flowers. To generate a final flower color segmented image, we combined these segmented images by applying a logical AND operation to the mask images, as illustrated in Figure 3. The estimated flower cover was quantified as the percentage of flower pixels relative to the entire image.

**Figure 3:**
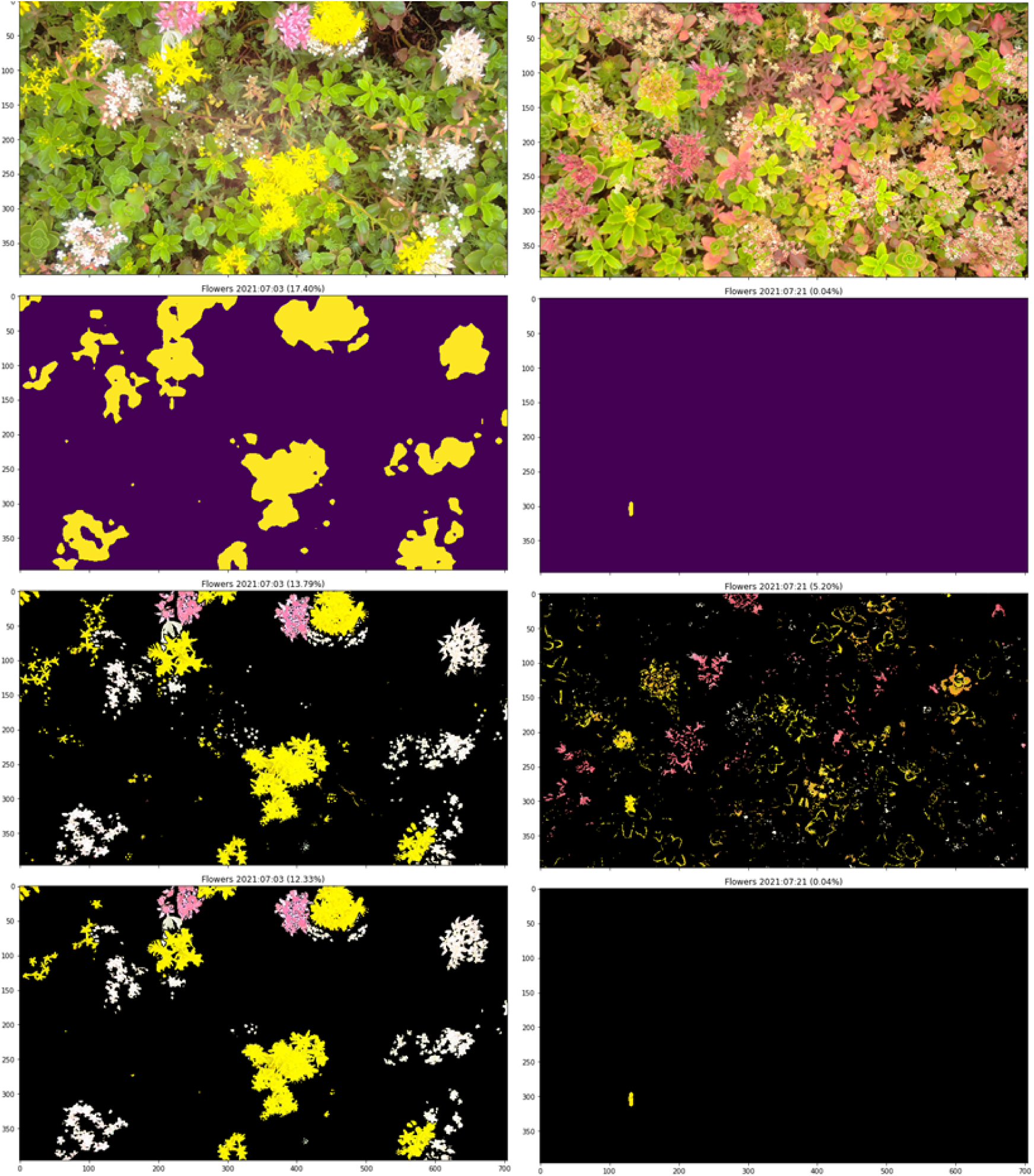
This figure presents examples of flower-segmented images. The top row displays the original images featuring Sedum plants. The second row shows the semantic segmentation results using DeepLab, highlighting the locations of the flowers. The third row illustrates the color-segmented images, identifying flowers of white, yellow, and pink. In the third row, the left image illustrates false flower segmentations. The bottom row combines the semantic and color-segmented images from the second and third rows, providing the final segmentation result with correct flower coverage.

### 2.2. Arthropod detection, filtering and classification

The image pipeline shown in Figure 1 consists of three stages, commencing with the detection (A) of arthropods, followed by filtering (B) of duplicate and false detections and concluding with the classification (C) of arthropods into taxonomic ranks.

While deep learning networks have the capability to integrate both detection and classification within the same model, we opted to divide the task into two distinct models. This approach offers increased flexibility, particularly because the specific taxa of arthropods present in the recordings are unknown until the detection phase is concluded. Additionally, training a generic detection model simplifies the annotation process, as classifier training can be conducted on grouped and sorted image crops of arthropods.

#### A Detection

Initially, arthropods were detected using a YOLOv5 (Redmon and Farhadi, 2018; Terven and Cordova-Esparza, 2023) model trained based on the methodology proposed in (Bjerge et al., 2023a,b). YOLOv5 provides the capability of models with different neural network architectures and image sizes. The smaller models allow faster execution. To improve performance and speed up training, the YOLOv5 is pre-trained on the COCO (Lin et al., 2015) dataset. We fine-tuned the pre-trained models and compared the capability of YOLOv5m6 to detect arthropods with a resized image size of 1280 *×* 768 pixels.

A dataset of annotated arthropods with motion-information-enhanced (MIE) images was created as described in Section 3.2. The MIE technique, as proposed by Bjerge et al. (2023b), extracts motion from the time-lapse sequence of images and combines it with color information to enhance the visibility of insects. The MIE technique was evaluated using state-of-the-art models, achieving the highest *F* 1 score with YOLOv5 compared to Faster-RCNN (Ren et al., 2017). The YOLOv5 detector trained on MIE images, resulting in an improved average *F* 1-score from 0.49 to 0.71 on a time-lapse recorded dataset containing insects in a similar setup to our studies. Our annotated dataset with various arthropods was used to fine-tune a pre-trained YOLOv5m6 model on the COCO (Lin et al., 2015) dataset with 35.7M parameters. The model was chosen because their size allows it to be run with the required memory and achieve high accuracy on most reasonably priced GPUs and achieved the best performance by (Bjerge et al., 2023a).

#### B Filtering

Detecting and localizing small arthropods within intricate and dynamic scenes of natural vegetation poses a challenge, especially when they may be occluded by flowers or leaves. The complexity lies in distinguishing the objects of interest from the natural vegetation. Despite the feasibility of achieving the detection and classification of arthropods *in situ*, the aforementioned challenges can cause models to erroneously identify background elements as objects of interest, particularly in the case of time-lapse images. A significant portion of these images may not contain any arthropods, resulting in false positive detections of plant parts in the background. To address this issue, especially given the temporal context of time-lapse images, we introduce a filtering mechanism. This filter named stationary aims to mitigate false detections by identifying and removing instances that occur in successive recorded images. It operates by disregarding detections found in the same position (distance less than 15 pixels) within the last 15 minutes, thereby reducing the likelihood of double-counting arthropods and eliminating some of the false positives.

In addition, we incorporated a matching filter designed to eliminate detections when no discernible difference is detected within the last three successive images of the recordings. This filter effectively eliminates detections of non-moving arthropods or false positives in the background.

The matching filter utilises the normalised sum of squared differences eq. (1) between the detected bounding box and the average of the corresponding detected bounding boxes in the preceding two recorded images eq. (2).

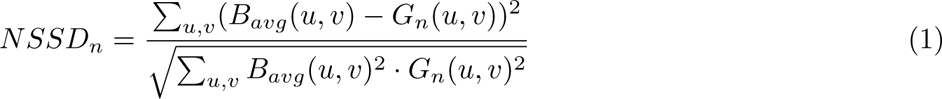

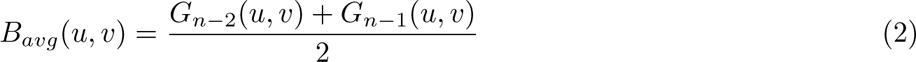

here, (*u, v*) represents the pixel coordinates within the bounding box of the detected object. *G*_*n*−1_ and *G*_*n*−2_ denote the detected bounding box in two preceding grayscale images. *B_avg_*signifies the average grayscale bounding box of the two preceding images. *G_n_* refers to the detected bounding box in the current grayscale image. The normalised sum of squared differences (*NSSD_n_*) provides a value indicating the similarity between the background bounding box *B_avg_* and the grayscale detected bounding box *G_n_*. A value of *NSSD_n_* close to zero indicates a match of the detection with preceding images and suggests that the detection may correspond to a non-moving arthropod or a false positive background detection. In such cases, the detection is flagged for elimination. An *NSSD_n_* threshold of 16 was empirical selected to match non-moving detections, based on samples from sequences containing moving arthropods and stationary detections. A match filtering with *NSSD* is illustrated in Figure 4.

**Figure 4:**
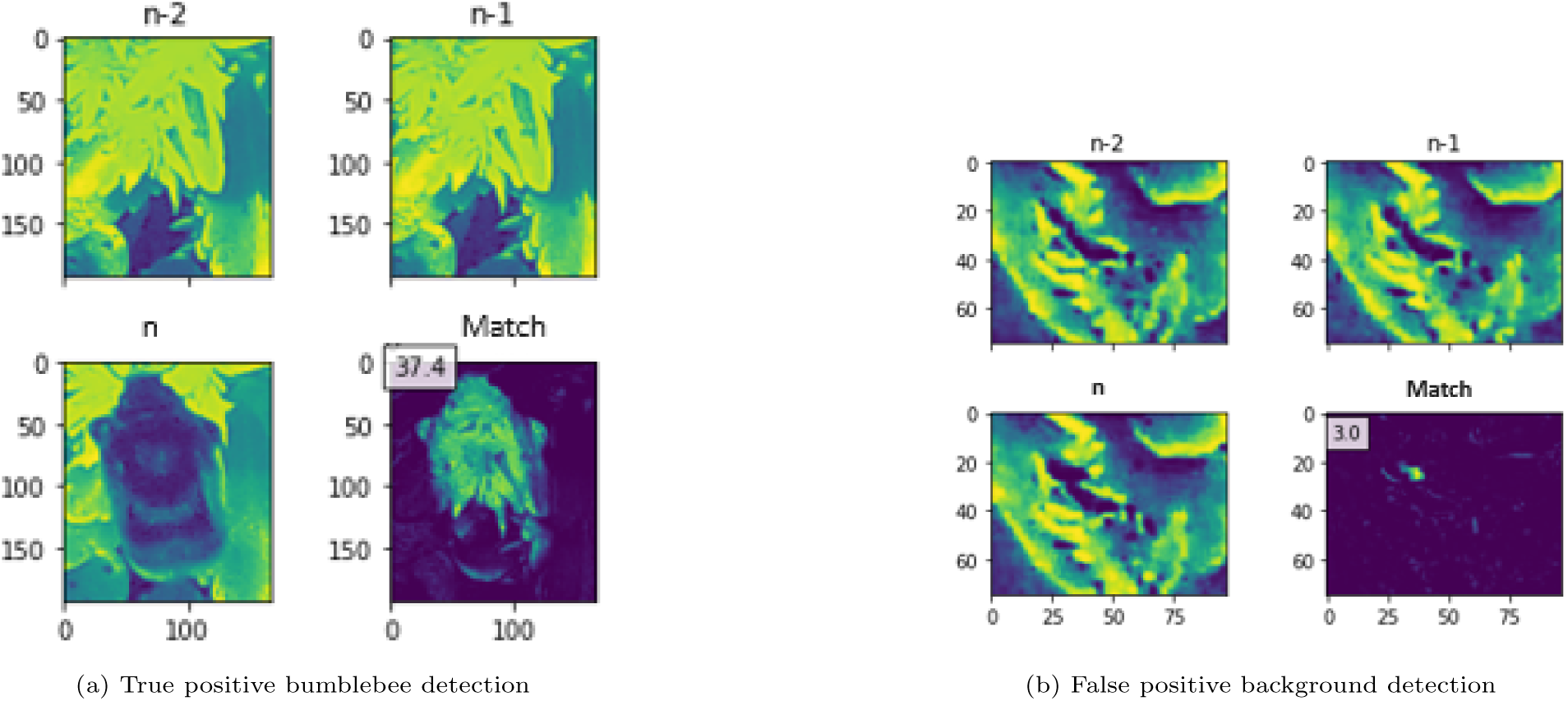
The left images shows three successive images of the bounding box for the detected bumblebee. With NSSD = 37.4 indicating a true positive object detection. The right images shows three successive images of the bounding box for a false positive detection. With NSSD = 3.0, the detection associated with an NSSD below 16 will be eliminated.

#### C Classification

In the classification problem within deep learning, the data is categorised into different classes, such as sorting arthropod images according to taxonomic ranks. The classes are defined by a number of different labelled categories in a dataset for training and test of the deep learning model. Here, we have sorted the detected and filtered arthropods in classes based on a manual study of the actually detected objects creating a dataset with 19 classes as described in Section 3. The dataset was partitioned into 80% for training and 20% for validation.

The training on the datasets was performed using data augmentation, including image scaling, horizontal and vertical flip and adding color jitter for brightness, contrast and saturation. Data augmentation mitigates overfitting by increasing the diversity of the training data. We selected a batch size of 32 for training our models since it is faster to update and results in less noise, than smaller batch sizes. The accuracy of the models on the training and validation datasets was computed after each epoch. The Adam optimizer with a fixed learning rate of 1.0 · 10^−4^ was finally chosen based on earlier experiments (Bjerge et al., 2023c; Smith, 2018)

We have selected and evaluated seven common state-of-the-art (SOTA) CNN models: ConvNext (Tiny and Base) (Todi et al., 2023), DenseNet121 (Huang et al., 2017), EfficientNetB4 (Tan and Le, 2019), InceptionV3 (Szegedy et al., 2016), MobileNetv2 (Sandler et al., 2018) and ResNet50v2 (He et al., 2016). The SOTA models were modified and trained using transfer learning with pre-trained weights from ImageNet (Russakovsky et al., 2015). Transfer learning involves training a model on data from a source domain *T_S_* = *P* (*y|X*) and then transferring it to a target domain *T_T_* = *P* (*y|X*) – typically with less data available. In this case, ImageNet contains 1,000 classes with 1,281,167 images for training and 50,000 for testing, while the arthropod dataset only contains 19 classes with 13,812 images. For all models, a fully connected layer with a dropout rate of 0.2 was added and trained on the dataset, followed by fine-tuning of all base CNN layers.

### 2.3. Performance metrics for testing

To evaluate model performance, the precision, recall and *F* 1-score metrics were chosen for evaluation the final detection, filtering and classification pipeline. These metrics are based on true positive (*TP*), false positive (*FP*) and false negative (*FN*) detections and classification. Recall and precision were used in conjunction to obtain a complete picture of the model’s ability to find all arthropods and classify them correctly. To balance precision and recall, we used the *F* 1-score. The *F* 1-score is calculated as the harmonic mean of precision and recall, given in eq. (5) and it provides a balance between the two metrics. It prioritises the importance of detecting and classifying the classes correctly.

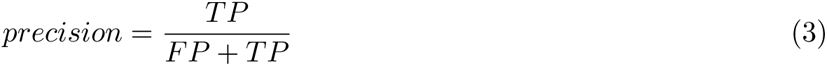

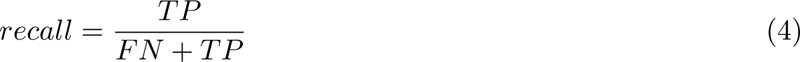

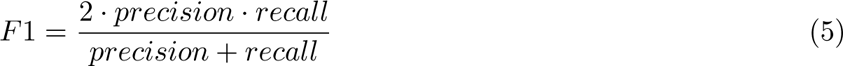

## 3. Experiment and datasets

Images were captured using Wingscapes (Wingscapes, 2024) cameras, which recorded time-lapse images at 60-second intervals from June to September in both 2020 and 2021, during the hours of 6 AM to 10 PM. The cameras were mounted 50 cm above the plants on custom-made metal frames, facing the ground. Images were captured at a resolution of 10 MP or 20 MP (4224 *×* 2376 or 6080 *×* 3420 pixels). We started the sampling with 20 MP, but reduced it to 10 MP as the Wingscapes camera internally upsize images, so increasing the resolution does not actually enhance image quality. The cameras were manually focused for close-range imaging.

The images encompassed *Sedum* plants, positioned at three distinct locations — urban, agriculture and grassland — each featuring four camera sites. All sites were located in Eastern Jutland, Denmark. The limited growth rate and height of *Sedum* plants of Sedum flowers ensure that the camera remains in focus throughout the entire monitoring season, enabling clear visible arthropods in the images. At each camera site, two plots measuring 120 *×* 80*cm* were cultivated with *Sedum* plants and monitored by two cameras. Hence, four closely located cameras were deployed at each location and site, resulting in a total of 3 *×* 4 *×* 4 = 48 cameras.

The primary objective was to showcase the feasibility of automated camera monitoring for live insects and flowers *in situ* on a comprehensive temporal and spatial scale. In total 5,932,597 images were recorded from the 48 cameras in the summer 2020 and total 4,524,328 images were recorded from the 48 cameras in the summer 2021. For model training and validation, only images from 2021 have been used.

### 3.1. Datasets with flower masks

Daily, at 12:00, an image was chosen from each of the 48 cameras in 2021, wherein a color mask, as detailed in Section 2.1, was applied to generate a binary mask for flowers in yellow, white and pink hues, as depicted in Figure 2 and 3. We also experimented with using images taken at 4:00 PM, but this did not result in any significant differences, as the background of the flower images remains relatively consistent throughout the day. Images were specifically selected if they exhibited more than 1% pixel area coverage of flowers, yielding 1,635 images featuring flower masks. Subsequently, images with false positive flower masks were eliminated, resulting in a refined dataset comprising 507 images with accurately segmented flowers. To augment the dataset, 25% of the images with false positive flower masks were reintroduced, with corrections made by a corrected black flower mask. The resultant dataset, featuring correct flower masks, encompasses a total of 768 images.

### 3.2. Datasets with arthropods

For insect detection and classification, all images recorded in 2021 were used to create two datasets: one for detection and one for classification, as described below. Despite the large full image resolution (4224 *×* 2375 or 6080 *×* 3420 pixels), each insect observation covered only a small portion of the full image, as the recording was conducted at a distance to monitor a designated patch of vegetation. Bjerge et al. (2023a) introduced a YOLOv5 model trained for insect detection on *Sedum* plants; however, this model only identified 26,987 detections in the recorded images equal to only 0.7% of the 3.9 million images. We utilised this model to generate a dataset comprising annotated bounding boxes for arthropods. The model yielded positive detections and false negative, encompassing instances of plant parts and animals, such as snails and birds. Subsequently, these detections of plant parts and animals were excluded and removed from the final dataset. A total of 6,901 images were meticulously selected, manually corrected and annotated with bounding boxes outlining arthropod locations. To enhance the detection of small objects, we employed motion-informed image enhancements, following the methodology proposed by Bjerge et al. (2023b). The resulting dataset, comprising annotated arthropods, is detailed in Table 1. Examples of background images and annotated arthropods are shown in Appendix A.

**Table 1:** Dataset used to train the detector of arthropods. It shows the number of images with number of annotated arthropods and background images without arthropods.

| Dataset | Images | Arthropods | Background |
| --- | --- | --- | --- |
| Training | 6,901 | 4,770 | 2654 |
| Testing | 766 | 525 | 299 |

A YOLOv5 model without MIE images was trained to detect arthropods and found 351,505 detections in image recordings from 2021. These detections were cropped and manually selected and sorted into 19 classes of taxonomic ranks as shown in Table 2. The images are semi-randomly chosen from the model detections trying to have as many images of individuals in each class. An example of sample images for each class is shown in Figure 5. The class of unspecified arthropods (Label ID 11) contains images of species with only few images insufficient to create separate classes. A class (Label ID 2) with false positive plant parts was also included in the group of classes. The resulting dataset is unbalanced ranging from 165 to 1842 images in each class. To facilitate model training, squared crops of arthropod images were resized to 224 *×* 224 pixels.

**Figure 5:**
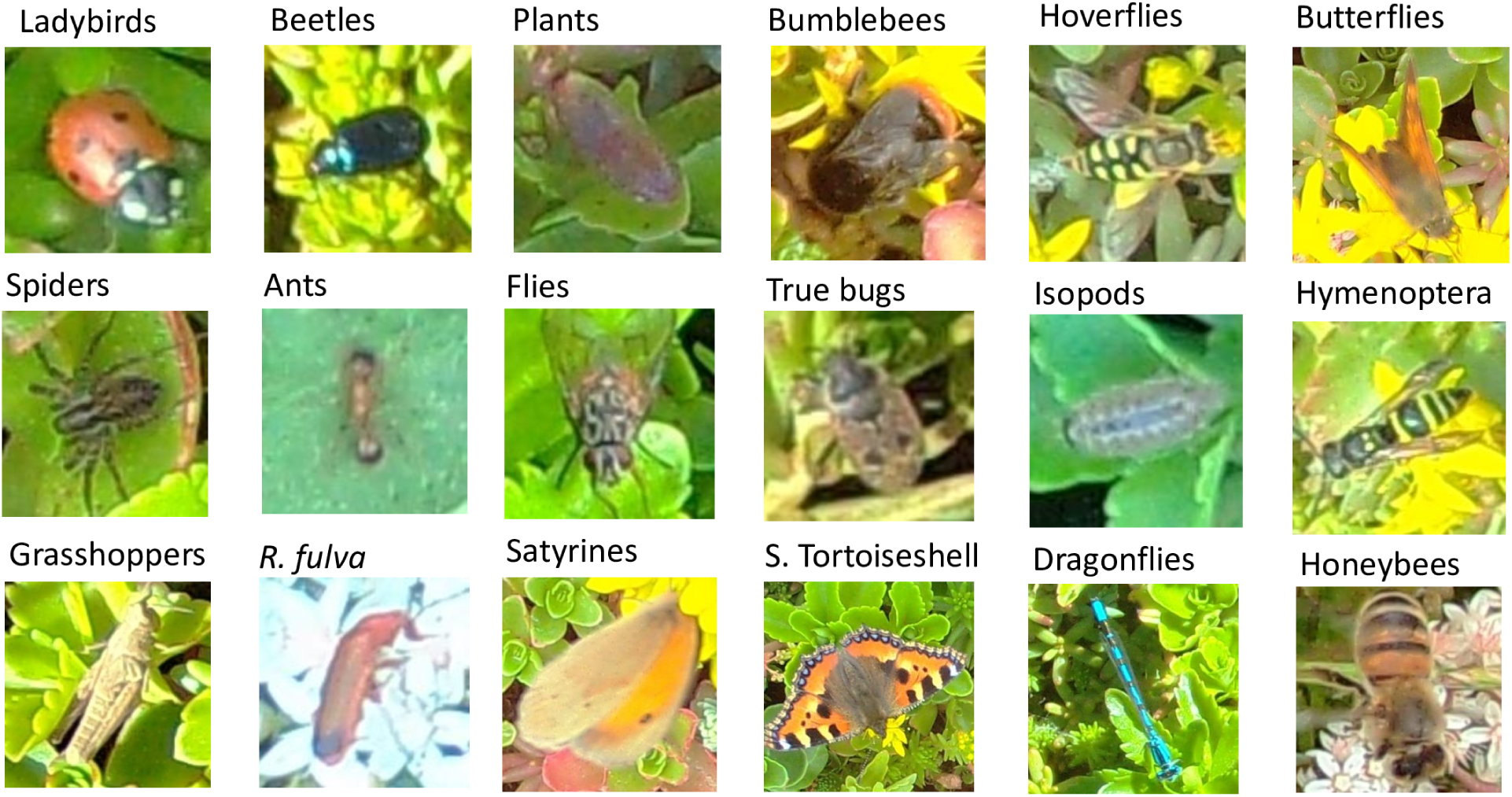
Example images of arthropods and plant parts for classes listed in Table 2

**Table 2:** The 19 classes of arthropods and plant parts used to train the classifier with in total 13,812 sample images. The classes of arthropods are grouped by different taxonomical ranks. The class ”Unspecified” is a mixture of different species with insufficient samples to create separate classes. The taxonomic rank of order, family, genus and species are listed for each class given by class name and a label ID. The splitting of the dataset is 20% for testing and 80% for training.

| Label ID | Class name | Order | Family | Genus/Species | Samples |
| --- | --- | --- | --- | --- | --- |
| 0 | Ladybirds | Coleoptera | Coccinellidae |  | 370 |
| 1 | Beetles | Coleoptera |  |  | 366 |
| 2 | Plants | Crassulaceae |  |  | 619 |
| 3 | Bumblebees | Hymenoptera | Apidae | Bombus | 616 |
| 4 | Hoverflies | Diptera | Syrphidae |  | 1,517 |
| 5 | Butterflies | Lepidoptera |  |  | 903 |
| 6 | Spiders | Aranaea |  |  | 1,842 |
| 7 | Ants | Hymenoptera | Formidicidae |  | 608 |
| 8 | Flies | Diptera |  |  | 663 |
| 9 | True bugs | Hemiptera |  |  | 638 |
| 10 | Isopods | Isopoda |  |  | 851 |
| 11 | Unspecified |  |  |  | 1,544 |
| 12 | Hymenoptera | Hymenoptera |  |  | 372 |
| 13 | Grasshoppers | Orthoptera |  |  | 165 |
| 14 | *R. fulva* | Coleoptera | Cerambycidae | *Rhagonycha fulva* | 820 |
| 15 | Satyrines | Lepidoptera | Nymphalidae | Satyrinae | 686 |
| 16 | Small tortoiseshell | Lepidoptera | Nymphalidae | *Aglais urticae* | 620 |
| 17 | Dragonflies | Odonata |  |  | 249 |
| 18 | Honeybees | Hymenoptera | Apidae | *Apis mellifera* | 363 |

## 4. Results and discussion

### 4.1. Estimating flowers in images

Utilizing the dataset outlined in Section 3.1, we trained a DeepLabv3 (Chen et al., 2018) semantic segmentation model. The optimal model was identified after 24 epochs out of 30 training epochs, demonstrating the lowest test loss (0.00675), an *F* 1-score of 0.505 and an Area Under Curve (AUC) of 0.888. The test and training losses converged to the same level, indicating an absence of overfitting. Subsequently, we combined the semantic-segmented images with flower color-segmented images through a logical AND operation on the mask images. The efficacy of this final model was assessed by randomly selecting 500 images from the 2020 and 2021 datasets. Among these, 15 images from 2020 and 151 images from 2021 exhibited false positive instances with flower segmentation.

Table 3 presents the percentage of flower pixel area for false positives (FP) and true positives (TP). Training the model on images from 2021 resulted in a notably high precision of 98.1%. However, when evaluating the model on images recorded in 2020, the precision drops to 80.7%. This discrepancy can be attributed to the fact that more images in 2020 contained plants with a different appearance than those in 2021. Example images can be found in Appendix A, where plants are overexposed or exhibit a more reddish hue, especially late in the season. This change is most certainly due to drought for much of the 2020 season that led to changes in the color profile of the plants. To enhance performance, the inclusion of images from both 2020 and 2021 in the training set for the semantic segmentation model could have been considered.

**Table 3:** Shows the false positive (FP) and true positive (TP) area of flowers in percentage of pixels out of 500 random selected images in 2020 and 2021. The precision of flower detection is calculated based on the FP and TP pixel areas.

| Year | FP pixel area<br>(%) | TP pixel area<br>(%) | Precision |
| --- | --- | --- | --- |
| 2020 | 223.0 | 932.6 | 0.807 |
| 2021 | 8.04 | 413.9 | 0.981 |

### 4.2. Detecting, filtering and classifying arthropods

A YOLOv5 model was trained on the detection dataset presented in Table 1, with and without motion-informed enhancement (MIE). The results in Table 4 demonstrate a high *F* 1-score for both models. The precision is 93.3% for the YOLOv5 model without MIE, compared to 89.0% for the model with MIE. In the paper by Bjerge et al. (2023b), the MIE method was proposed to increase model performance on new recordings. The study found that when a dataset is split into training and validation sets, the MIE method does enhance performance on data from new unseen locations despite reduced validation performance. This improvement demonstrates the MIE’s generalization ability and its robustness against overfitting. As a result, the MIE method proves valuable when testing on datasets recorded at different locations than those used for training and validation. The model trained with MIE showed in general a higher number of detections than the model trained with color images, indicating more insects were found and detected on a time-lapse test dataset created from different sites and periods of the growing season not included in the training and validation dataset.

**Table 4:** Shows the precision, recall, *F* 1-score and mean Average Precision (mAP) of the YOLOv5 models trained without and with motion-informed enhancement (MIE).

| Model | Precision | Recall | *F1*-score | mAP |
| --- | --- | --- | --- | --- |
| YOLOv5 | 0.933 | 0.901 | 0.917 | 0.943 |
| MIE | 0.890 | 0.908 | 0.908 | 0.935 |

While the precision of arthropod detection is quite high at 90%, this still results in a considerable number of false positive predictions when dealing with millions of time-lapse images. The magnitude of this issue is illustrated in Table 5, where a substantial number of false positive plant detections are observed. These are subsequently filtered using a sequence of stationary and matching filters applied after the YOLOv5 MIE arthropod detector.

**Table 5:**
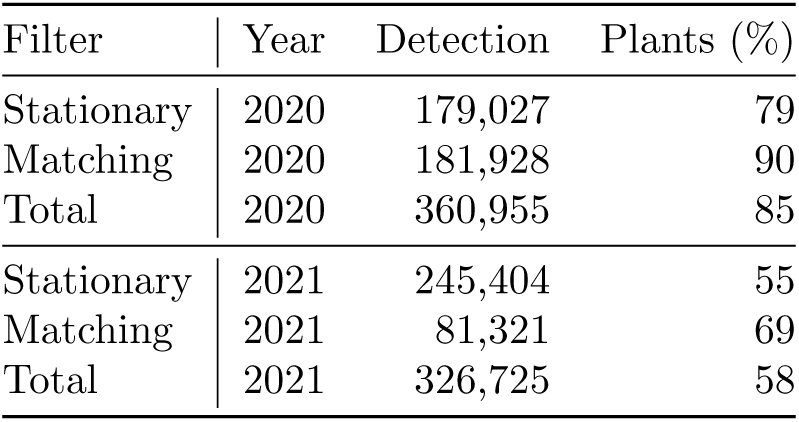
Shows the filtered detections by the stationary and matching filters of the image recordings in 2020 and 2021. The percentages of plants (and other elements in the background) are evaluated on 200 randomly selected detections from both 2020 and 2021. Here the remaining percentages of detections were identified as arthropods. The total number of detections filtered by the stationary and matching filters are summarized with the percentage of plants.

The stationary detections encompass both false plant detections and non-moving arthropods, aiming to minimize the double counting of the same individual. However, if an animal has moved within the camera’s field of view, it may be counted multiple times. The matching filter further mitigates false positive plant detections, particularly in 2020, where 90% of the 181,928 detections pertain to plant parts. Notably, detections filtered by the matching filter may include arthropods that blend with the background, resulting in their removal.

The purpose of the next processing pipeline step was to sort the filtered detections in classes of arthropod taxa. The dataset with 19 classes of arthropods was used to train and fine-tune the seven state-of-the-art CNN models, as shown in Table 6. The results indicate that fine-tuning all layers of the CNNs improved performance, with the *F* 1-score increasing from 0.81 to 0.85 for EfficientNetB4. The best performance was achieved by ConvNeXt-Base, which attained an *F* 1-score of 0.89. However, this model is resource-intensive, with a significant number of parameters and requires more than 25GB of GPU memory for training. EfficientNetB4 ranks among the top three performing models while having significantly fewer parameters compared to the ConvNeXt family of CNN models. EfficientNetB4 was finally selected as the model for classifying arthropods in our pipeline due to its strong performance relative to its computational complexity.

**Table 6:** Summary of state-of-the-art CNN models fine-tuned and validated on 20% of the dataset. The table presents the number of model parameters (in millions), along with the precision, recall, and *F* 1-score metrics, all evaluated on the validation dataset with 19 classes of arthropods.

| Model | Parameters<br>millions | Precision<br>validation | Recall<br>validation | *F1*-score<br>validation |
| --- | --- | --- | --- | --- |
| MobileNetV2 (Sandler et al., 2018) | 6.779 | 0.816 | 0.811 | 0.813 |
| ResNet50V2 (He et al., 2016) | 72.192 | 0.834 | 0.834 | 0.834 |
| InceptionV3 (Szegedy et al., 2016) | 65.456 | 0.850 | 0.845 | 0.847 |
| DenseNet201 (Huang et al., 2017) | 54.617 | 0.849 | 0.847 | 0.848 |
| EfficientNetB4 (Tan and Le, 2019) | 52.873 | 0.853 | 0.847 | 0.850 |
| ConvNeXt-Tiny (Todi et al., 2023) | 83.504 | 0.858 | 0.842 | 0.850 |
| ConvNeXt-Base (Todi et al., 2023) | 262.757 | 0.891 | 0.885 | 0.888 |

The selected EfficientNetB4 model was trained on the arthropod dataset, which consists of 19 different classes, as detailed in Section 3.2. Analysing the confusion matrix in Figure 6, it becomes evident that certain classes, namely plants (2), butterflies (5), true bugs (9), unspecified (11) and Hymenoptera (12), exhibit a subpar accuracy, falling below 70%. The low accuracy for plants (2) is attributed to YOLOv5 arthropod detections that could not be effectively filtered by the stationary and matching filter. The low accuracy in identifying butterflies (5) stems from numerous images featuring butterflies with folded wings, posing a challenge for classification. The unspecified (11) class comprises a mix of arthropods that could not definitively be grouped into any other class. Consequently, many arthropods are erroneously classified into this unspecified category, such as 49% of Hymenoptera (12). True bugs (9) exhibit low accuracy due to misclassifications as various other classes, including beetles, plants, spiders, grasshoppers and unspecified categories. The precision, recall and *F* 1-score for each class are provided in more detail in Table A.1.

**Figure 6:**
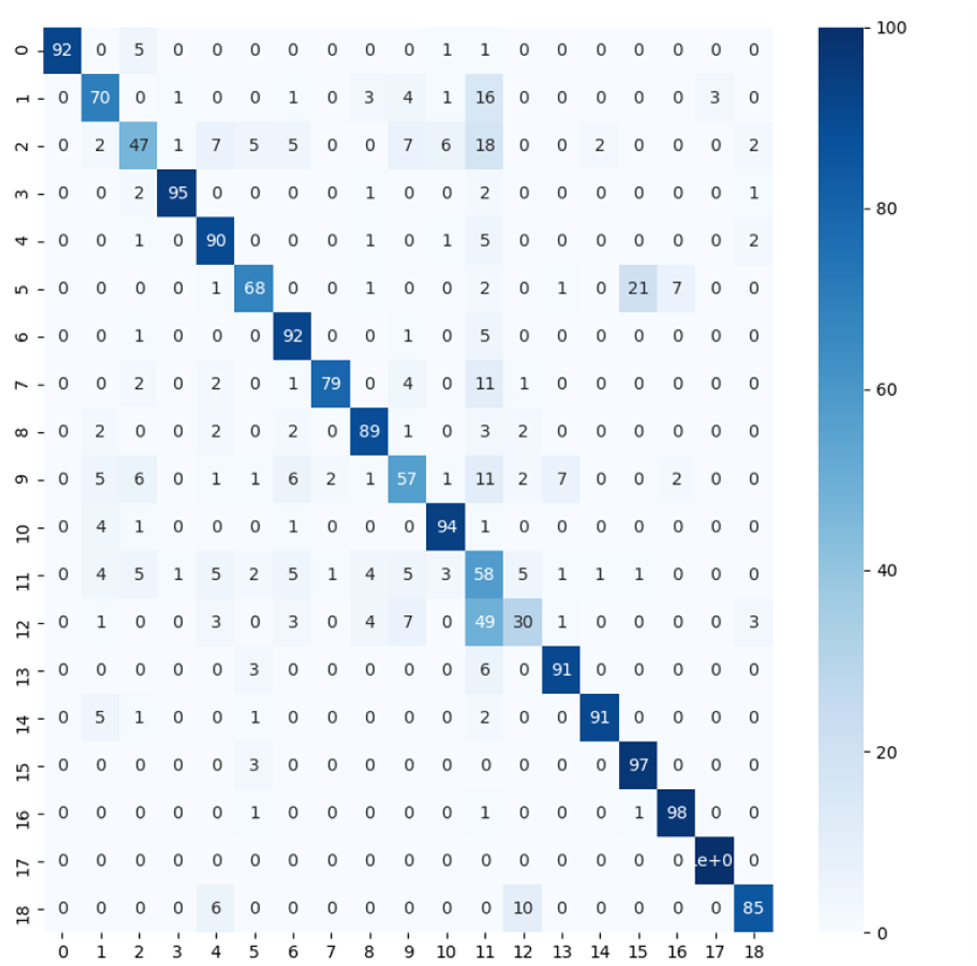
Confusion matrix on the validation dataset of the trained EfficientNetB4 model. The X-axis is the ID of the predicted labels. The Y-axis is the ID of the true labels. The numbers in the table are the accuracy percentage related to true labels. A high diagonal value indicates a high accuracy.

After the detection and filtering of the 2020 and 2021 recordings, predictions were generated using the trained EfficientNetB4 model. The count of predictions for each of the 19 classes is presented in Table 7. A validation test was conducted on 200 randomly selected predictions from each class. Class predictions that could not be confidently verified were manually labeled as false positives. For the unspecified class, only plant parts were treated as false positive predictions, resulting in a higher precision than that achieved on the validation dataset in Table A.1. Given that the EfficientNetB4 model was trained on data recorded in 2021, a higher precision of 80% was attained for data from 2021 compared to the 60% precision in 2020. Classes with the best performance, exhibiting a precision above 60%, include ladybirds (0), plants (2), hoverflies (4), spiders (6), ants (7), flies (8), Hymenoptera (12), grasshoppers (13) and R. fulva (14). Notably, butterflies (5), bumblebees (3), true bugs (9), satyrines (15) and dragonflies (17) show precisions below 30% in 2020. In the analysis of 200 random test samples, it is particularly plant parts that are misclassified in the mentioned classes.

**Table 7:** Shows the number of classified predictions of 19 categories of arthropods in 2020 and 2021. Precision 2020/2021 is a test performed on 200 random predictions selected from each class after detection and filtering on the entire image collection in 2020 and 2021. The total number of predictions and the average precision in 2020 and 2021 are listed at the bottom of the table. The best performing classes with a precision above 60% in 2020/2021 are marked with an ’*’.

| Label ID | Class name | Predictions<br>2020 | Predictions<br>2021 | Precision<br>2020 | Precision<br>2021 |
| --- | --- | --- | --- | --- | --- |
| 0 | Ladybirds* | 556 | 580 | 0.64 | 0.88 |
| 1 | Beetles | 5,478 | 4,813 | 0.58 | 0.70 |
| 2 | Plants* | 34,537 | 19,027 | 0.90 | 0.92 |
| 3 | Bumblebees | 1,216 | 1,634 | 0.35 | 0.80 |
| 4 | Hoverflies* | 15,437 | 6,620 | 0.90 | 0.82 |
| 5 | Butterflies | 5,749 | 3,218 | 0.07 | 0.56 |
| 6 | Spiders* | 16,048 | 23,862 | 0.72 | 0.80 |
| 7 | Ants* | 2,867 | 6,412 | 0.92 | 0.94 |
| 8 | Flies* | 8,708 | 13,162 | 0.98 | 0.98 |
| 9 | True bugs | 6,207 | 8,946 | 0.29 | 0.46 |
| 10 | Isopods* | 8,708 | 4,380 | 0.80 | 0.80 |
| 11 | Unspecified* | 68,351 | 38,664 | 0.78 | 0.90 |
| 12 | Hymenoptera* | 411 | 2,556 | 0.78 | 0.88 |
| 13 | Grasshoppers* | 2,320 | 1,462 | 0.76 | 0.70 |
| 14 | *R. fulva* * | 7,949 | 3,708 | 0.66 | 0.74 |
| 15 | Satyrines | 2,303 | 3,173 | 0.25 | 0.80 |
| 16 | Small tortoiseshell | 158 | 691 | 0.50 | 0.98 |
| 17 | Dragonflies | 385 | 285 | 0.07 | 0.81 |
| 18 | Honeybees | 964 | 509 | 0.48 | 0.70 |
|  | Total/Average | 187,866 | 143,702 | 0.60 | 0.80 |

In the years 2020 and 2021, we identified approximately 300,000 arthropods among the more than 10 million recorded images. The inherent challenge with time-lapse recordings in our scenario lies in the relatively low occurrence of arthropods compared to the vast number of images, leading to a significant number of false positive plant detections compared to the number of positive arthropod detections. With a precision of 89% measured on the validation dataset in table 4 there should in theory be 11% false positive detections which will result in more than 1 million false plant detections in the recordings from 2020 and 2021. If we combine the estimates of plant detections in table 5 and table 7 an approximately 550,000^1^ false plant detections. Therefore, the filtering approach we propose is crucial for mitigating this issue by reducing the number of false positive plant detections before classification into different taxa of arthropods.

### 4.3. Evaluation of flower estimation and arthropod detection and classification

Figure 7 illustrates the temporal progression of the results for a designated camera positioned in a grassland environment throughout the year 2021. At the onset and conclusion of the season, characterised by diminished flower coverage, there is a noticeable absence of insects. The prevalence of spiders mirrors the abundance of other arthropods, consistent with their role as predators. Notably, the utilization of YOLOv5 with MIE leads to the detection of a greater number of small arthropods, including spiders and ants. The same tendency was observed by Bjerge et al. (2023b) where the models trained MIE achieved a high F1-score on a time-lapse test dataset.

**Figure 7:**
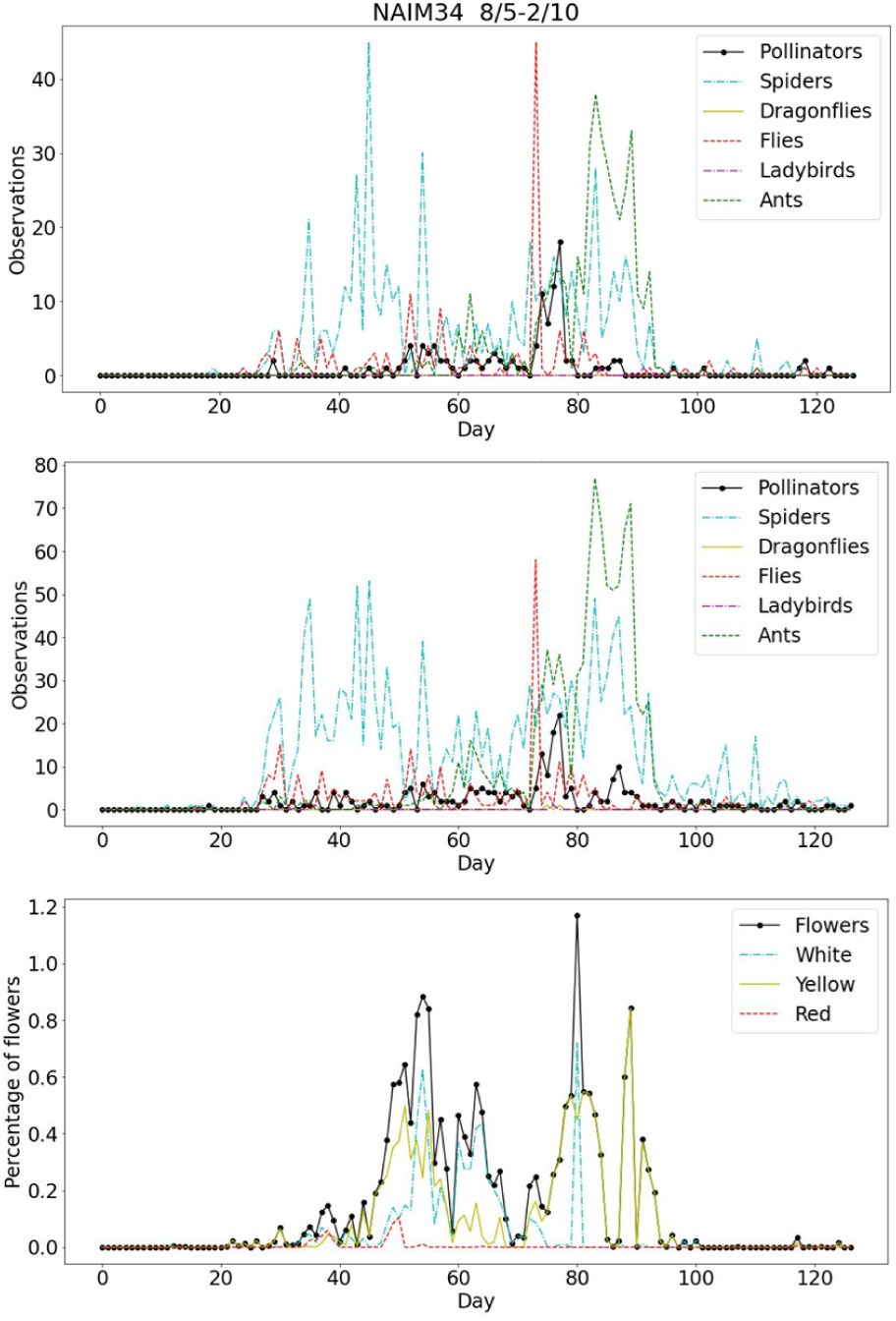
The analytical outcomes derived from a particular camera (NAIM34) situated in a grassland setting during the year 2021. The upper image illustrates the outcome of the arthropod detection, filtering and classification pipeline using YOLOv5 without MIE for a specific set of classes. Pollinators encompass bumblebees, hoverflies, honeybees and butterflies (Satyrines and S. tortoiseshell). The middle image displays the pipeline results with YOLOv5 and MIE. The lower image exhibits the flower cover, highlighting white, yellow and red flowers.

Figure 8 depicts the temporal dynamics observed in two cameras situated within agricultural and grassland settings throughout the year 2021. The figures show distinct increase in abundance of pollinators during the flowering phase. This consistent trend is corroborated by two additional camera locations in the urban habitats, as depicted in fig. 8b. Notably, spiders, certain flies, isopods, and ants show activity even outside the flowering period. While some flies and likely all spiders do not visit flowers for nectar or pollen, they may still act as pollinators, either as predators or while basking. The presence of spiders and other insect pests on plants is known to influence pollinator behaviour, sometimes through plant-mediated effects, resulting in complex changes in pollinator dynamics Knauer et al. (2018). Our method provides a valuable tool for gaining deeper insights into these intricate pollinator interactions.

**Figure 8:**
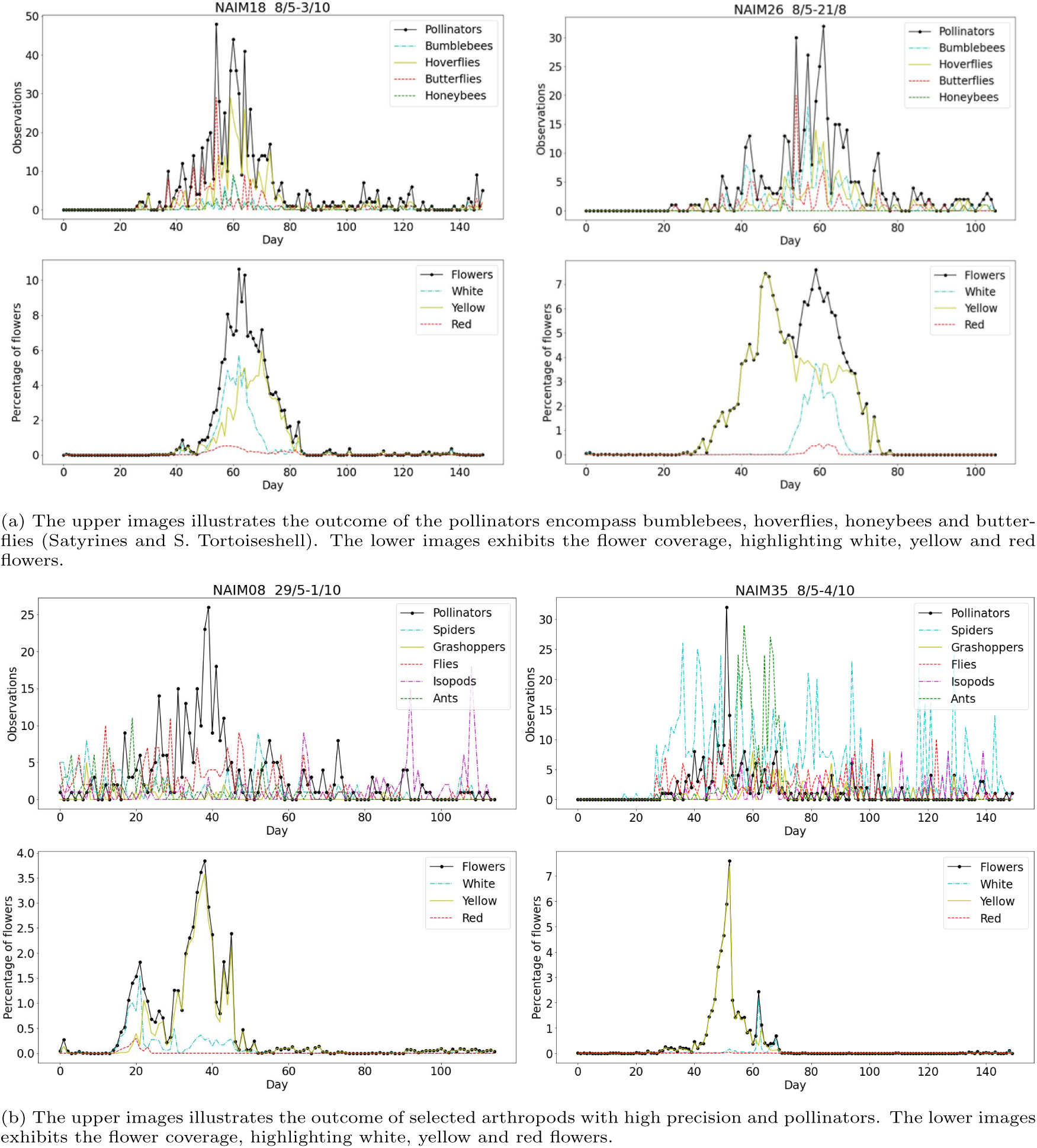
Shows the analytical outcomes derived from four cameras situated in an urban (NAIM08), agriculture (NAIM18) and grassland (NAIM26, NAIM35) setting during the year 2021.

### 4.4. Discussion on limitations and potentials

We observed that the abundance of pollinators at the beginning and end of the season is reduced, mirroring the decrease in flower coverage during these periods. The high correlation between the abundance of pollinators and flower cover indicates that cameras placed above plants with abundant flowers are more likely to record active insects. Consequently, in order to compare insect abundance across sites, it is important to correct for variation in flower cover. Ideally this should be done at the plant species level. It may also be relevant to relocate cameras during the season to follow flowering patterns. This could increase the number of recorded insects.

During the experiments, some data was lost for instance due to camera batteries dying and corruption of SD cards. Using cameras with embedded computers connected to the internet, as employed in Bjerge et al. (2021a), could mitigate manual data loss. However, building and deploying such cameras require more technical skills and may be more expensive compared to the cost-effective Wingscapes cameras used in our experiments. Monitoring insects in natural vegetation is challenging due to difficulties with camera focusing, especially with plants of varying heights. This is why we chose low-growing *Sedum* plants for our study. The camera trap design by Sittinger et al. (2024) includes high frame-rate tracking of insects. To address the camera focusing issue these insect observations are made against an artificial background, rather than in a natural environment of flowers and plants. Extending real-time video monitoring, as conducted by Ratnayake et al. (2021); Bjerge et al. (2021a), to include flower cover estimation, as proposed in our paper, could provide more comprehensive studies of insects in natural wildlife environments.

While our research has focused on DeepLabv3, many other state-of-the-art DL semantic segmentation models could also be explored. However, our findings suggest that an immediate improvement would be to extend the dataset by including flower masks from both years, thereby increasing the number and diversity of images in the training set.

Unlike *Sedum*, many other plants have complex structures that insects navigate, and their flowers often appear against varied backgrounds such as leaf litter, rocks, sticks, bark, and other natural objects, potentially reducing the effectiveness of color masks. Further research would be needed to adapt our method into a general flower estimator.

The detected and classified arthropods are assigned to one of the 19 taxonomic groups that the model has been trained to recognize. Consequently, any species not included in the model’s training set will be incorrectly classified into one of these 19 categories. To address this limitation, the model could be enhanced with a hierarchical classifier incorporating anomaly detection, as proposed by Bjerge et al. (2023c). This approach would allow for the classification of uncertain or new insect species at a higher rank within the taxonomy hierarchy.

The AMI dataset published by (Jain et al., 2024) was used to evaluate five state-of-the-art models for nocturnal insect species classification. Among these, ConvNext-Base (Todi et al., 2023) and ViT-B/16 (Dosovitskiy et al., 2021) were identified as top performers and could be considered as alternatives to the Efficient-NetB4 model we employed in our work. On the AMI-GBIF dataset, ConvNext-Base achieved an accuracy of 90.45%, outperforming ResNet50, which had an accuracy of 87.56%. On the AMI-Traps dataset, ViT/B achieved an accuracy of 83.01%, outperforming ResNet50, which had an accuracy of 78.95% on insect species. We have observed a similar increase in accuracy with ConvNext-Base compared to ResNet50 in our study. However, our results suggest that further improvements could be achieved by curating and enhancing our dataset, specifically by incorporating images from both years of our experiment. This would increase the diversity and number of sample images available for training the arthropod classifier.

Existing manual methods for studying and estimating the composition of plant communities, such as those described by Tabatabai (1998), are labour-intensive. Camera-based methods, as used in other studies by Körschens et al. (2024), have shown that automatically predicting species-wise plant cover closely reflects original estimates made by human experts at the same sites. We anticipate that a similar conclusion will be reached for flower coverage, with automated estimation from images likely to be accurate and more efficient than manual methods.

Our approach holds immediate applicability to image-based insect monitoring in natural settings, serving purposes in agriculture and biodiversity conservation (Ratnayake et al., 2021; Bjerge et al., 2021a; Preti et al., 2021). We have showcased its capability in estimating flower cover and establishing a discernible correlation between pollinator abundance and recorded flower cover. Our method can generate relevant information regarding flower phenology by analyzing color variations over time and species abundance as studied by Mann et al. (2022). This allows for the exploration of insect preferences towards different flower species.

## 5. Conclusion and future work

In conclusion, our proposed methodology presents a comprehensive pipeline for estimating flower cover and detecting arthropods in natural flowering environments. The utilization of semantic and color segmentation provides a straightforward approach to estimating flower cover in images. However, it is only demonstrated on a single variety of *Sedum* plants with colors of white, yellow and red/pink.

Given the relatively low arthropod occurrence within complex vegetation, our solution addresses the impact of false positives on detection precision. The incorporation of stationary and matching filters effectively eliminates false detections of arthropods and reduces the occurrence of double-counting individual arthropods. Over half of the detections involve false positives or non-moving animals, underscoring the importance of our filtering mechanisms. The final step of the pipeline is a trained CNN classifier that categorizes arthropods into selected taxa based on our experiment’s actual recordings.

Our comprehensive evaluation, conducted across 48 cameras in three distinct habitats with over 10 million images, resulted in datasets used to train a YOLOv5 object detector and an EfficientNetB4 classifier. The pipeline exhibited an average precision of 80%, with certain categories, such as hoverflies, ants, and flies, achieving precision levels exceeding 90%. Furthermore, our study confirmed a strong correlation between insect pollinators and flowering plants.

Future efforts should focus on collecting additional time-lapse images with annotated arthropods to train a generic YOLO detector using motion-informed enhanced images for broader applicability across monitoring sites. Further enhancing the taxonomic resolution of the arthropod classifier, as further data is collected, is realistic and highly relevant. This would also allow for grouping taxa into functional groups and likely also increase the performance of the classifier. Moreover, replacing the unspecified arthropods class with an anomaly detector, achieved by thresholding CNN classifier output scores, could further refine the classification process. An ecological analysis of the results is left to future studies outside the scope of this paper.

## CRediT authorship contribution statement

Kim Bjerge: Conceptualization, Data curation, Investigation, Methodology, Software, Validation, Visualization, Writing – original draft. Hjalte M.R. Mann: Data curation, Writing – review & editing. Toke T. Hoye: Conceptualization, Data curation, Validation, Funding acquisition, Project administration, Writing – review & editing. Henrik Karstoft: Supervision, Writing – review & editing.

## Acknowledgments

The work was supported by the European Union’s Horizon Europe Research and Innovation programme, under Grant Agreement No. 101060639 (MAMBO). K.B., H.M.R.M. and T.T.H. were also funded by the EU Horizon 2020 Research and Innovation programme [Grant Agreement no. 773554 (EcoStack)]. We would like to give thanks to the companies Nature Impact and Byggros that have provide *Sedum* plants for all 48 camera setups.

## Data availability

The source code for the deep learning pipeline and monitoring results (arthropods abundance and flower cover estimates from 2020 and 2021) can be downloaded from: https://github.com/kimbjerge/ insectsFlowers

## Declaration of Generative AI and AI-assisted technologies in the writing process

During the preparation of this work the author(s) used ChatGPT in order to improve formulations in writing. After using this tool/service, the author(s) reviewed and edited the content as needed and take(s) full responsibility for the content of the publication.

## Appendix A. Supplementary material

In Figure A.1, full-sized (6080 *×* 3420 pixels) images are shown with examples of background images without annotations. Examples of annotations from part of full-sized images are shown in Figure A.2. Examples of false positive flower segmentations are shown in Figure A.3 from images recorded in 2020. The precision, recall, and *F* 1-score of the EfficientNetB4 model, evaluated on 20% of the arthropod dataset without fine-tuning the CNN base layers, are shown in Table A.1. After fine-tuning the base layers, the *F* 1-score improved to 0.85, as shown in Table 6.

**Figure A.1:**
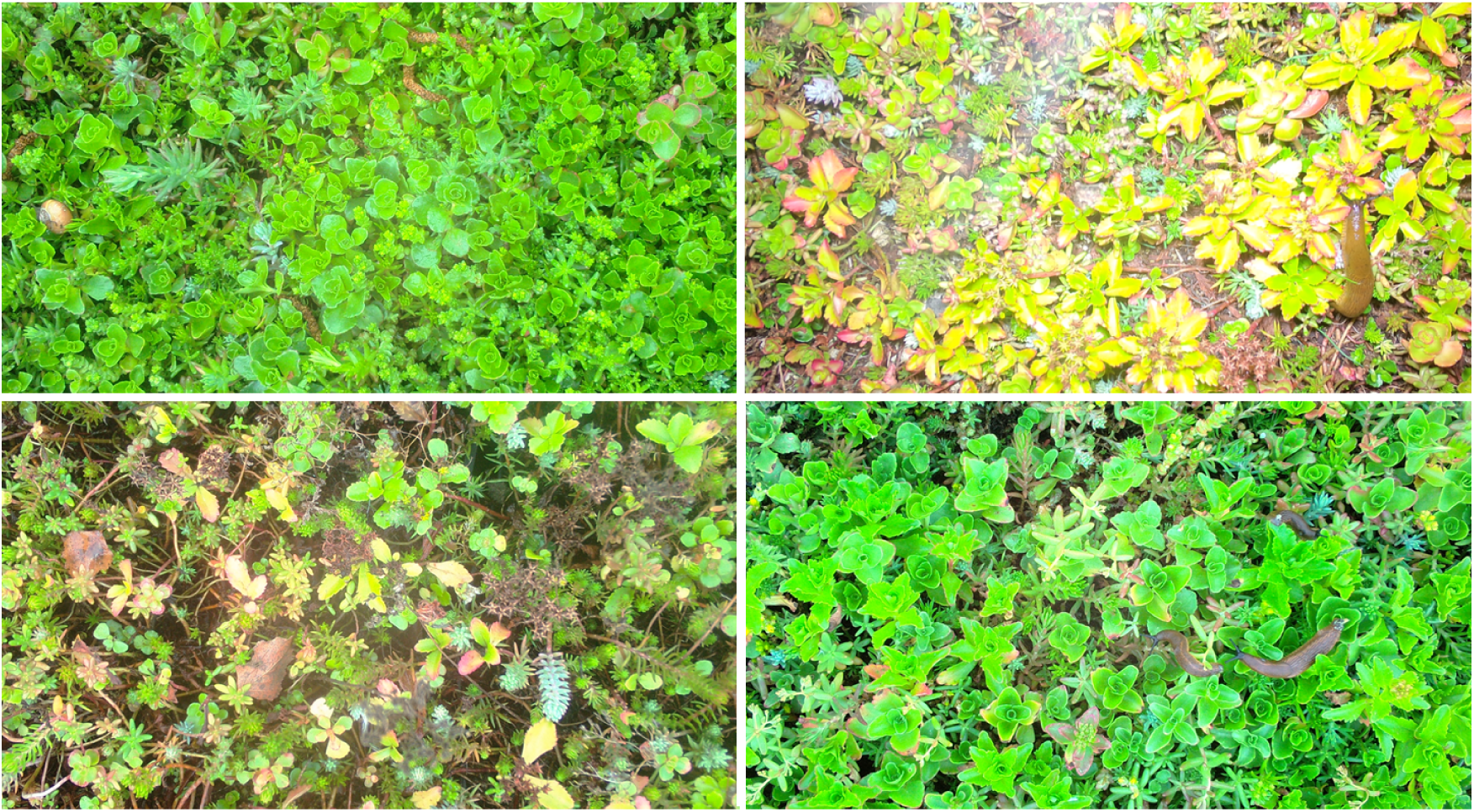
Full size images of background images with killer slugs (*Arion lusitanicus*) but without annotations of arthropods.

**Figure A.2:**
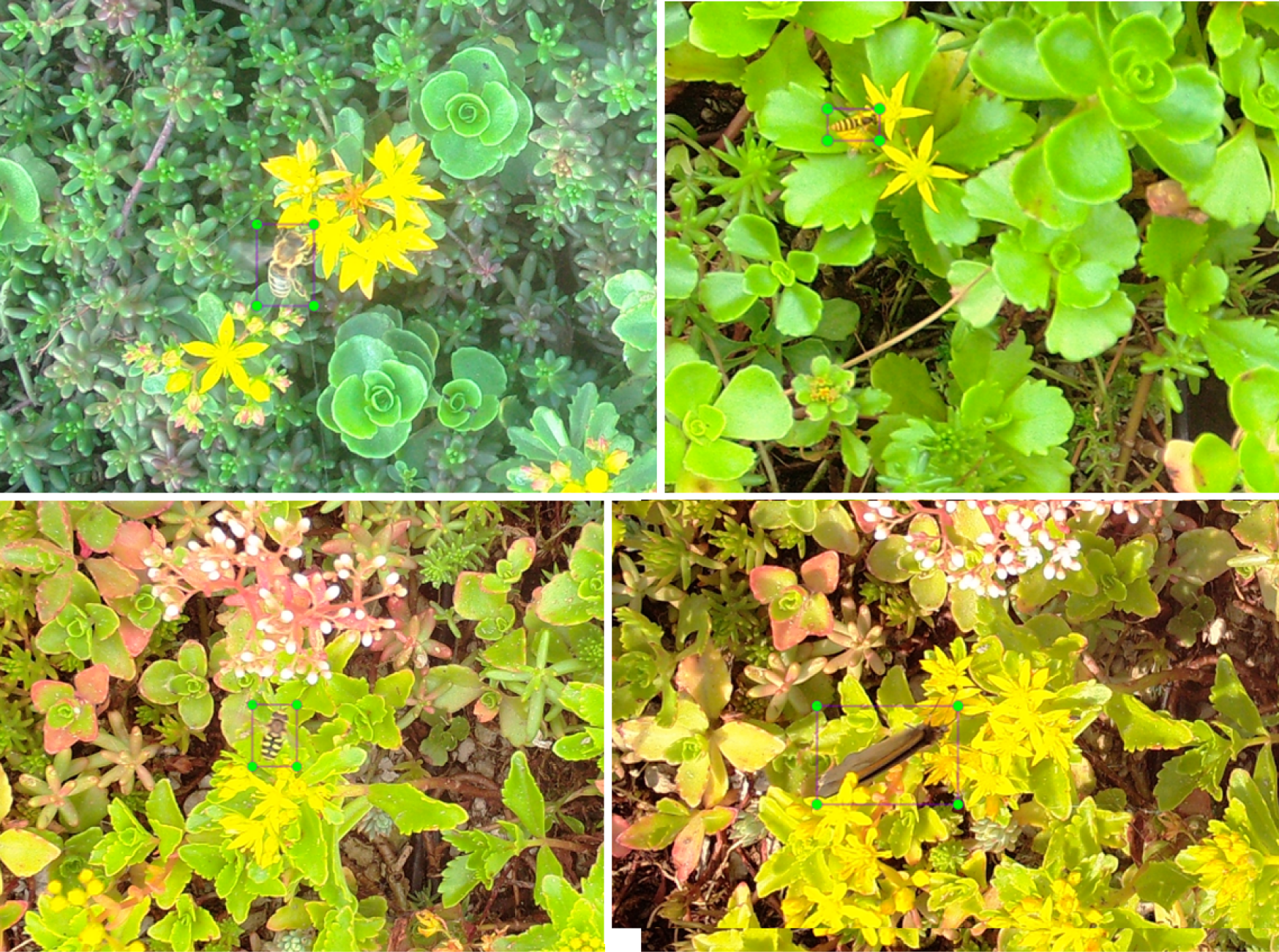
Part of full-size images with annotated arthropods (Honeybee, hoverflies and butterfly).

**Figure A.3:**
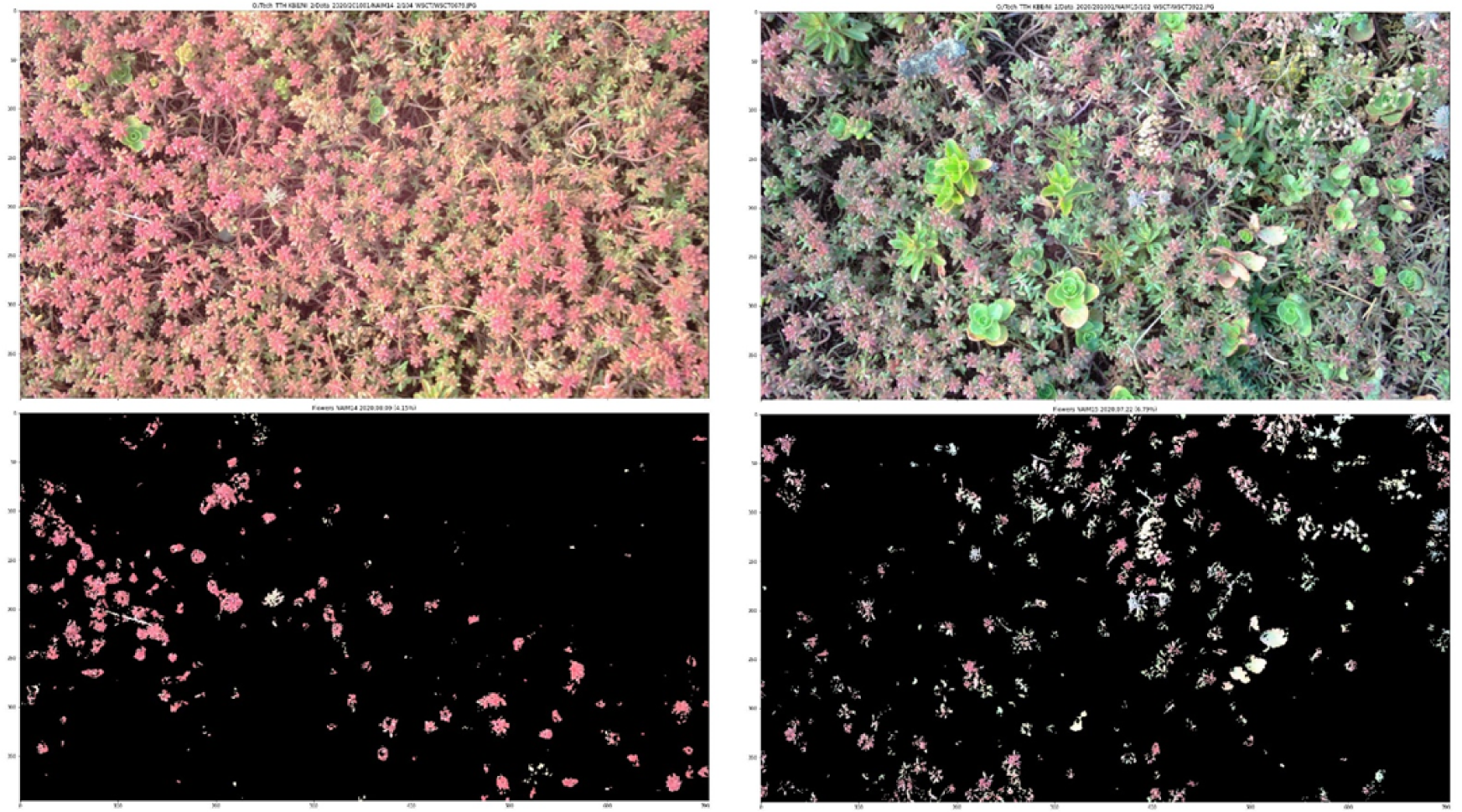
The top row displays original images. The second row shows example of false positive combined flower segmentations in images recorded in 2020.

**Table A.1:** The used EfficientNetB4 model validated on 20% of the dataset with the 19 classes of arthropods and plant parts in Table 2. The precision, recall and *F* 1-score are show evaluated on the validation dataset without fine-tuning of CNN base layers.

| Label ID | Class name | Precision<br>validation | Recall<br>validation | $F1$-score<br>validation | Validation<br>dataset |
| --- | --- | --- | --- | --- | --- |
| 0 | Ladybirds | 0.97 | 0.92 | 0.94 | 74 |
| 1 | Beetles | 0.57 | 0.70 | 0.63 | 73 |
| 2 | Plants | 0.60 | 0.47 | 0.53 | 123 |
| 3 | Bumblebees | 0.95 | 0.95 | 0.95 | 123 |
| 4 | Hoverflies | 0.88 | 0.90 | 0.89 | 303 |
| 5 | Butterflies | 0.87 | 0.68 | 0.76 | 180 |
| 6 | Spiders | 0.90 | 0.92 | 0.91 | 368 |
| 7 | Ants | 0.95 | 0.79 | 0.86 | 121 |
| 8 | Flies | 0.84 | 0.89 | 0.86 | 132 |
| 9 | True bugs | 0.64 | 0.57 | 0.60 | 127 |
| 10 | Isopods | 0.88 | 0.94 | 0.91 | 170 |
| 11 | Unspecified | 0.55 | 0.58 | 0.57 | 308 |
| 12 | Hymenoptera | 0.43 | 0.30 | 0.35 | 74 |
| 13 | Grasshoppers | 0.67 | 0.91 | 0.77 | 33 |
| 14 | *R. fulva* | 0.96 | 0.91 | 0.94 | 164 |
| 15 | Satyrines | 0.76 | 0.97 | 0.85 | 137 |
| 16 | Small Tortoiseshell | 0.88 | 0.98 | 0.93 | 124 |
| 17 | Dragonflies | 0.96 | 1.00 | 0.98 | 49 |
| 18 | Honeybees | 0.82 | 0.85 | 0.84 | 72 |
|  | Average | 0.80 | 0.81 | 0.81 |  |

## Footnotes

1 360, 955 × 0.85 + 34, 537 × 0.90 in 2020, 326, 725 × 0.58 + 19, 027 × 0.92 in 2021

